# Decoding the Influence of Anticipatory States on Visual Perception in the Presence of Temporal Distractors

**DOI:** 10.1101/143123

**Authors:** Freek Van Ede, Sammi R Chekroud, Mark G Stokes, Anna C Nobre

## Abstract

While it has long been recognised that anticipatory states amplify early EEG responses to visual targets in humans, it remains unclear how such modulations relate to the actual content of the neural representation, and help prioritise targets among temporally competing distractor stimuli. Using multivariate orientation decoding of high temporal resolution EEG recordings, we first demonstrate that anticipation also increases the amount of stimulus-identity information contained in these early brain responses. By characterising the influence of temporally adjacent distractors on target identity decoding, we additionally reveal that anticipation does not just attenuate distractor interference on target representations but, instead, delay it. Enhanced target decoding and distractor resistance are further predicted by the attenuation of posterior 8-14 Hz alpha oscillations. These findings offer several novel insights into how anticipatory states shape neural representations in service of resolving sensory competition in time, and they highlight the potential of non-invasive multivariate electrophysiology to track cognitive influences on perception in tasks with rapidly changing displays.

**Highlights:**

- Anticipatory states help resolve visual competition in time
- Anticipation enhances early target coding and delays distractor interference
- Attenuated alpha oscillations also enhance target coding and distractor resistance
- EEG decoding is a powerful tool for tracking percepts in rapidly changing displays

**Significance statement:** While the neural mechanisms by which anticipatory states help prioritise inputs that compete in space have received ample scientific investigation, the mechanisms by which the human brain accomplishes such prioritisation for inputs that compete in time remain less well understood. We used high temporal resolution EEG decoding to individuate (and track in time) neural information linked to visual target and distractors stimuli that were presented in close temporal proximity. This revealed that anticipatory states help resolve temporally competing percepts by a combination of enhanced target (but not distractor) coding as well as delayed interference on this target coding caused by temporally adjacent distractors – thus allocating a “protective temporal window” for high-fidelity target processing.

## Introduction

In a world in which the amount of information that reaches our senses is increasing by the day, it is becoming increasingly relevant to understand the mechanisms by which our brains extract and prioritise information that is most relevant to current goals. Foreknowledge of what, where or when relevant events are likely to occur enables the instantiation of anticipatory neural states that provide key determinants of such prioritisation (Posner, 1980; Nobre et al., 2011), and it has long been recognised that such anticipatory states amplify early brain responses to perceptual targets. In fact, such effects provided the first clear evidence in humans that modulatory effects of anticipatory attention occur early during sensory processing (e.g., Mangun and Hillyard, 1987, 1991; Luck et al., 1994). Yet, despite a long tradition, vast literature, and sustained interest in this line of research (for reviews, see e.g., Hillyard & Anllo-Vento, 1998; Luck et al., 2000; Eimer, 2014), it has remained unclear whether anticipation actually amplifies the amount of information defining the identity of the perceptual target in these early EEG responses. Building on recent progress on multivariate decoding of visual orientation information from high temporal resolution M/EEG measurements (e.g. Ramkumar, 2013; Garcia, 2013; Myers, 2015; Cichy, 2015; King and Dehaene, 2016; see also Stokes et al., 2015), we tackled this issue directly and reveal that anticipatory states also amplify stimulus-identity information.

Multivariate decoding with high temporal resolution additionally enabled us to individuate neural information linked to target vs. competing distractor items occurring within the temporal window of attentional competition. While the neural mechanisms that prioritise inputs that compete in space have received ample scientific investigation (for reviews, see e.g., Desimone and Duncan, 1995; Kastner and Ungerleider, 2000; Squire et al., 2013; Anton-Erxleben and Carrasco, 2013), the mechanisms by which the human brain accomplishes such prioritisation for inputs that compete in time remains far less well understood. This is in part because conventional human neuroimaging approaches have been hampered either by insufficient temporal resolution (as with fMRI), or by the presence of strong additive responses when stimuli occur in fast temporal succession (as with classical ERP analyses). By combining stimulus orientation decoding analyses with high temporal resolution EEG measurements, we reveal that anticipatory states not only enhance neuronal target representations, but also delay the interference caused by temporally adjacent distractors, thereby providing an extended protected temporal window for target analysis.

## Results

### Task and EEG orientation decoding

Thirty healthy human volunteers performed a visual orientation reproduction task in which the presence/absence of preparatory auditory cues and temporally adjacent visual distractors were orthogonally manipulated (Fig. 1a; Methods for details). Auditory cues, when present, indicated the target would follow after 500 ms, thus acting as temporal warning signals. Provided that our main research questions regard largely unexplored territory, we deliberately focused on an experimental design with such simple (but highly effective) temporal warning cues. While we will refer to the influence of these cues as anticipation, we acknowledge up front that this type of anticipation likely involves a mix of involuntary increases in vigilance and voluntary orienting of attention in time (Weinbach and Henik, 2012, for further discussion).

**Figure 1.**
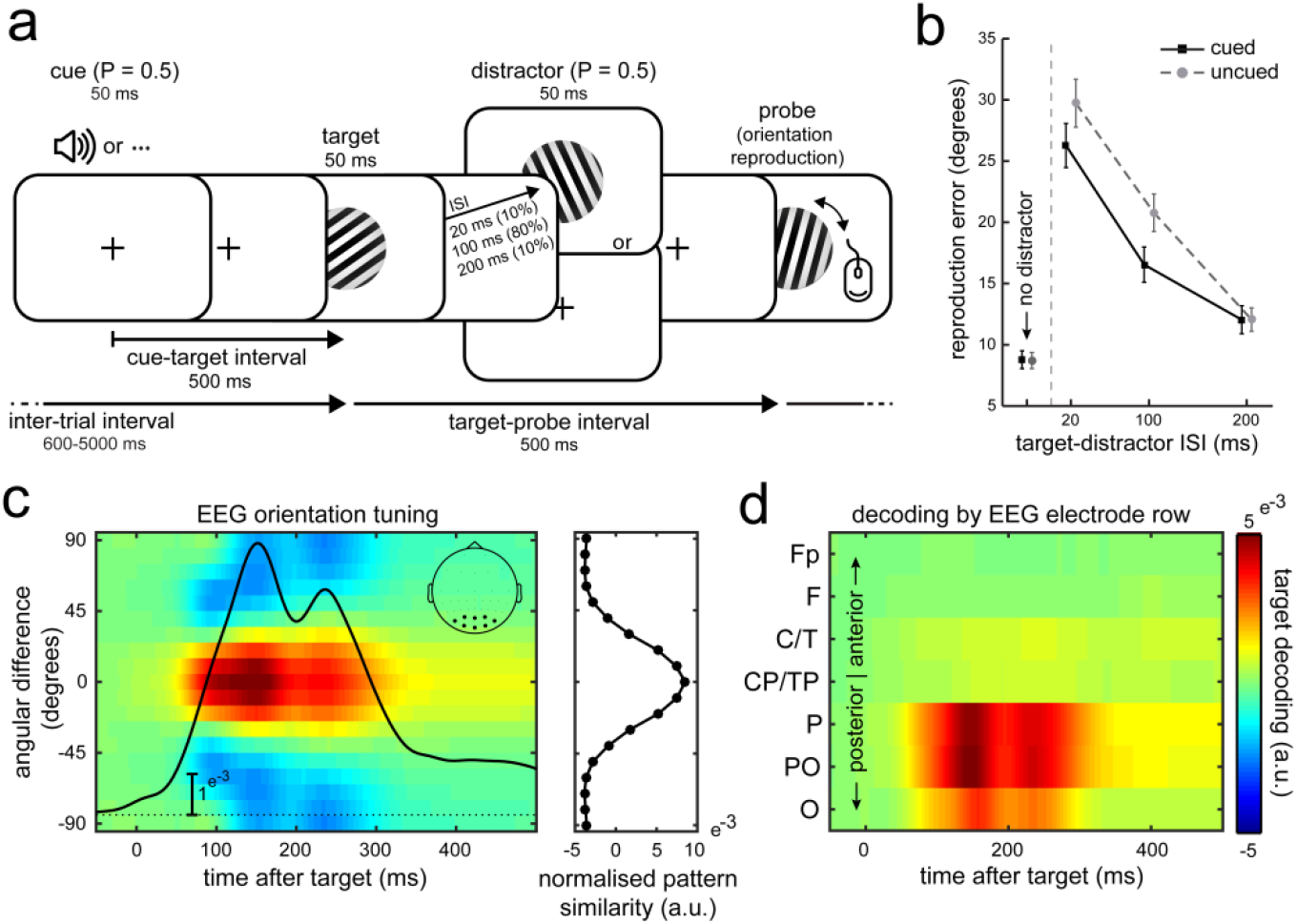
Task design, perceptual performance, and EEG orienting decoding. **(a)** Visual orientation reproduction task with preparatory auditory cues and visual distractors. Participants reproduced the orientation of the visual target grating using a computer mouse. In half the trials, targets were preceded by an auditory warning cue. Targets could be followed by no distractors, or by a visual distractor at one of three ISIs (20, 100, 200 ms). Target-probe intervals and inter-trial intervals were drawn independently of cue and distractor presence. **(b)** Average orientation reproduction errors (in degrees) for cued and uncued trials as a function of distractor presence and ISI. Error bars represent ± 1 SE calculated across participants (n = 30). **(c)** Time resolved orientation tuning profile based on all targets. Data represent the mean normalised pattern similarity (quantified using the Mahalanobis distance) between the test trials and the training trials, as a function of the angular difference between test and training trials (y axis). The inset highlights the 8 electrodes that were used for the orientation decoding analysis. Overlaid timecourse depicts the associated summary decoding statistic (Methods for details). The right panel shows the associated tuning profile averaged over the interval between 0 and 500 ms post-target. **(d)** Time resolved decoding (summary statistic) as a function the EEG electrode row used for decoding.

Figure 1b depicts average reproduction errors and highlights the utility of the cue in reducing distractor interference. While we found no cueing benefit on performance in distractor-absent trials (*t*_(29)_ = 0.345, *p* = 0.733, *d* = 0.063), reliable cueing benefits occurred in distractor-present trials (i.e. lower reproduction errors to cued vs. uncued targets), which interacted with inter-stimulus-interval (ISI; *F*_(2,58)_ = 12.926, *p* = 2.277e-5, *η_p_^2^* = 0.781). Planned comparisons confirmed a moderate cueing benefit at the 20-ms ISI (*t*_(29)_ = -3.537, *p* = 0.001, *d* = -0.646), a large benefit at the 100-ms ISI (*t*_(29)_ = -6.476, p = 4.353^e-7^, *d* = -1.182), but no longer any benefit when distractors followed targets at an ISI of 200 ms (*t*_(29)_ = -0.02, *p* = 0.984, *d* = -0.004). Cueing benefits were also significantly larger in distractor-present compared to distractor-absent trials, both at 20-ms ISI (*t*_(29)_ = -3.617, *p* = 0.001, *d* = -0.66) and at 100-ms ISI (*t*_(29)_ = -7.291, *p* = 4.97e^-8^, *d* = -1.331). Because we had anticipated (based on prior piloting) that the 100-ms ISI would be particularly effective, we deliberately used this ISI in the vast majority (80%) of distractor-present trials and focused our EEG analyses exclusively on this set.

Our main aim was to investigate the influence of the preparatory cues and temporally competing distractors on the amount of sensory information contained in the EEG responses regarding the identity of target and distractor stimuli (i.e., grating orientation). To this end, we applied a time-resolved “decoding” approach. Per time point, we calculated the multivariate *Mahalanobis* distance (using electrodes as dimensions) between the left-out trial (the “test trial”) and all other trials (the “training trials”) in which the target orientation was at a particular angular difference from the test trial. By evaluating this multivariate distance metric for a range of angular differences between test and training trials, we were able to reconstruct an orientation tuning profile (as in Wolff et al., 2017).

Figure 1c illustrates the utility of this approach, by depicting the time-resolved tuning profile averaged over all target presentations (i.e. independent of experimental condition). Target evoked EEG responses are most similar to other targets that have similar orientations (red), relative to other target that have more dissimilar orientations (blue). Robust tuning was most evident between 75 to 300 ms after target onset. To capture this orientation decoding in single metric (per time point), we simply multiplied the (mean normalised) tuning profiles with an inverted cosine function and averaged the result along all angular differences between test and training trials (as in Sprague et al., 2016; Wolff et al., 2017). To illustrate that this summary statistic captures the EEG orientation tuning well, we superimposed its time course on the orientation tuning profile in Figure 1c. We report on this summary statistic in all further analyses.

To concentrate our decoding analysis on visual activity, we limited the decoding analysis to data from the eight most posterior electrodes (inset Fig. 1c), which also showed the largest ERP (Fig. S1a). To further substantiate the visual origin of the orientation decoding, we additionally ran this analysis separately for each of the electrode rows. As shown in Figure 1d, this confirmed a predominantly posterior (putatively visual) origin.

### Anticipation increases target decoding and delays distractor interference

We next evaluated EEG orientation decoding as a function of cue and distractor presence, and considered six (non-mutually exclusive) scenarios by which anticipatory states may help extract and prioritise relevant over irrelevant sensory inputs that compete in time (Fig. 2). As we detail below, we found evidence in support of scenarios 1 (enhanced target decoding) and 6 (delayed distractor interference).

**Figure 2.**
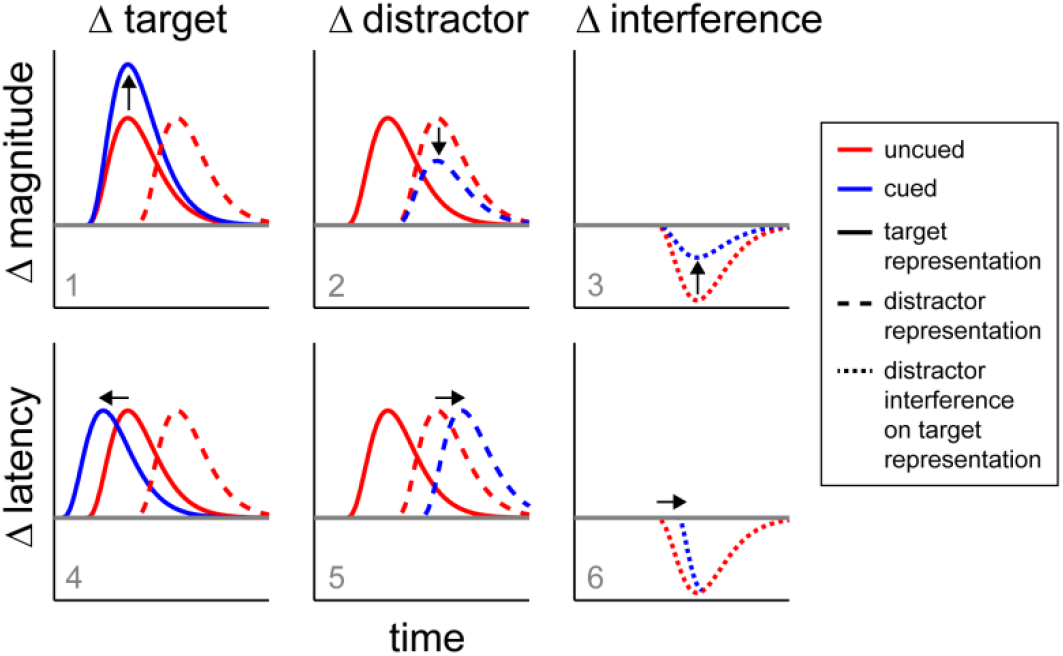
Schematic of ways in which anticipation may prioritise relevant over irrelevant sensory inputs that compete in time. We consider six non-mutually exclusive scenarios. Anticipation can influence the target representation (left), the distractor representation (middle), or the interference on the target representation when distractors are present compared to absent (right). This influence can be manifest either as a change in magnitude (“representation strength”), or a change in latency (“representation timing”). We find evidence for scenarios 1 (increased target identity decoding) and 6 (delayed distractor interference on the target identity decoding). See also Figure S2.

Figure 3a depicts time-resolved orientation decoding for each of the experimental conditions, both for targets and for distractors. Cluster-based permutation statistics (Maris and Oostenveld, 2007) revealed three significant clusters (corrected for multiple comparisons along the time axes) that each involved differences in target decoding (see also Fig. 3c). First, we observed a main effect of cue presence (cyan), as reflected in better orientation decoding for cued compared to uncued targets (cluster *p* = 0.006, cluster interval: 118 to 248 ms post target). This is in line with scenario 1 in Figure 2. In contrast, we found no significant cueing effect on distractor decoding (if anything, we observed a numerical increase, rather than a decrease, arguing against scenario 2 in Fig. 2). Second, we observed a main effect of distractor presence (magenta), as reflected in reduced target decoding for distractor present compared to distractor absent trials (i.e. distractor interference; cluster *p* = 0.004, cluster interval: 262 to 414 ms post target). Finally, we observed an interaction between cue and distractor presence (green; cluster *p* = 0.03, cluster interval: 196 to 268 ms post target). This effect was constituted by a larger cueing benefit for distractor present trials, or, equivalently, a larger distractor interference for cue-absent trials. While we note that the main effect of distractor presence on target decoding was maximal in the time window in which distractor decoding itself was also maximal, the interaction effect on target decoding was maximal in the time window in which the distractor decoding emerged (Fig. 3a). All three effects were again largely confined to the posterior electrode rows (Fig. 3c).

**Figure 3.**
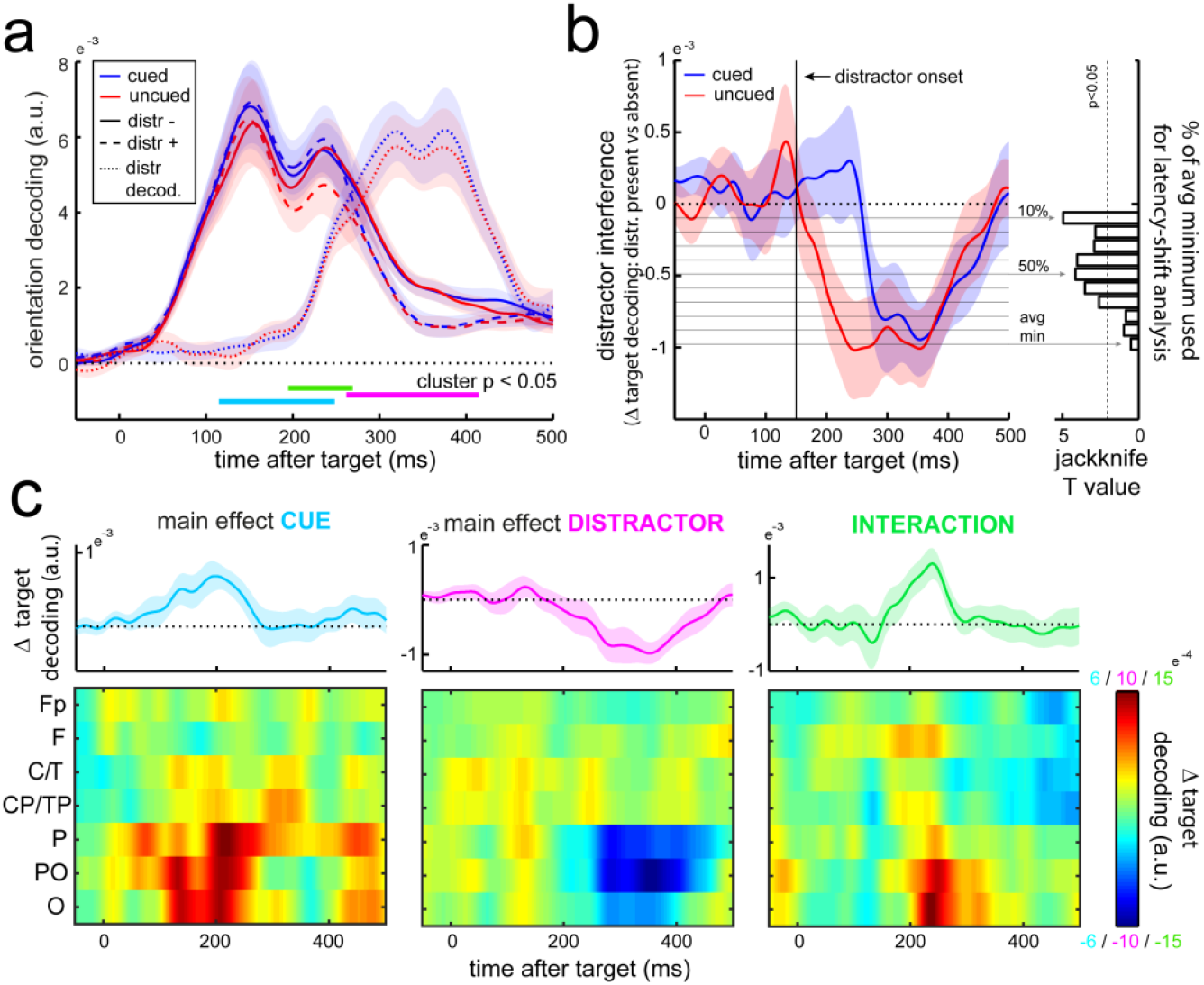
Anticipation increases target identity information and delays distractor interference in early visual EEG responses. **(a)** Time courses of target and distractor orientation decoding (summary statistic) as a function of cue and distractor presence. Horizontal lines indicate significant temporal clusters (Methods for details) for the main effects of cue presence (cyan), distractor presence (magenta), as well as their interaction (green). All clusters involve target decoding; no significant cueing effect cluster was observed for distractor decoding. See Figure S1 for the corresponding ERP results. **(b)** Time courses of the distractor interference effect on target decoding. Distractor interference is quantified as the difference in target decoding for distractor present vs. absent trials. Right panel shows Jackknife T values for latency differences between cued and uncued trials at thresholds ranging from 10 to 100% of the maximal interference effect. Maximal interference was calculated as the lowest value in the average of the cued and the uncued trials (denoted “avg min”). **(c)** Main and interaction effects as a function of time and electrode row. The interaction is expressed as the difference between cue-present vs. absent trials in distractor-present vs. absent trials. Upper plots show decoding based on the same channels as in **a** (see Fig. 1c). Shadings represent ± 1 SE calculated across participants (n = 30).

Figure 3b shows the time courses of the distractor interference effect on target decoding (i.e., target decoding with distractor present minus absent), and suggests that the observed interaction may be best understood as a delayed distractor interference effect (scenario 6 in Fig. 2). While both cued and uncued trials ultimately reach a similar level of distractor interference (unlike scenario 3 in Fig. 2), the onset of this interference appears delayed in cued trials. To further quantify this delay, we estimated the latencies at which the cued and uncued interference effects first reached the value associated with 50 percent of the maximal interference value (averaged over both conditions), and used a Jackknife approach (as described in Miller et al., 1998) to evaluate this delay statistically. This confirmed a 77 ± 18.57 ms (mean ± SE) delay in cued compared to uncued trials (Jackknife *t*_(29)_ = -4.145, *p* = 6.74e^-5^). Moreover, although we initially selected 50 percent of the maximum interference value for these analyses, it is reassuring to note that similar statistics were obtained when estimating latencies from values ranging anywhere from 10 to 70 percent of the maximum interference value (right panel Fig. 3b).

In contrast to scenarios 4 and 5 in Figure 2, Figure 3a showed no evidence for a cueing effect on the latencies of either target or distractor decoding alone (target: Jackknife *t*_(29)_ = 0.251, *p* = 0.299; distractor: Jackknife *t*_(29)_ = 0.342, *p* = 0.316). To provide further support against these scenarios, we also ran a cross-temporal decoding analysis whereby we trained the model on uncued trials and tested decoding performance on cued trials. Decoding was always best when train and test times corresponded (Fig. S2), thus providing further evidence that the EEG “orientation code” does not appear to shift forward (for targets) or backward (for distractors) in time with cueing.

Because our decoding was based on the broadband visual responses, a natural question is whether the observed cueing effects on target decoding and distractor resistance may simply be carried over from amplified ERP responses in cued trials (for example, by virtue of higher signal-to-noise ratio; SNR). When evaluating ERP amplitudes (Fig. S1a,c), we did also observe a main effect of cue presence that occurred at a similar time window as the main cueing effect on decoding (putatively reflecting amplification of the classic N1 potential). Interestingly, however, across our pool of 30 participants, the magnitude of this cueing effect on the ERP was uncorrelated with the magnitude of the cueing effect on decoding (*r* = -0.093, *p* = 0.624; Fig. S1d). In further contrast to the decoding results, we also did not observe an interaction between cue and distractor presence on the ERP that could account for the increased distractor resistance observed in decoding (Fig. S1b,c).

### *Attenuated posterior alpha oscillations predict enhanced target decoding and* distractor resistance

A key marker of attentional orienting in human M/EEG measurements is provided by the anticipatory attenuation of 8-14 Hz alpha oscillations in relevant sensory brain areas (Foxe et al., 1998; Worden et al., 2000, Thut et al., 2006; Mazaheri and Jensen, 2010; Foxe and Snyder, 2011; van Ede et al., 2012). Here we link such brain states in posterior electrodes to increased target decoding as well as distractor resistance, thereby also corroborating (using orthogonal analyses) the above described influences of the anticipatory cues.

Figure 4a shows the time- and frequency-resolved difference in spectral power between cued and uncued trials, averaged over all posterior electrodes. Immediately after the cue, we observed a transient increase in low-frequency power with a frontal-central topography (left inset Fig. 4a) that likely reflects cue processing. At a later stage, however, we also observed a decrease in 8-14 Hz power with a posterior topography (right inset Fig. 4a). Rather than cue processing, the latter likely reflects the instantiation of an “attentional brain state”. This state appears to emerge before target onset (in line with above references) although in our data it becomes most prominent during target and distractor processing (likely as consequence of the relatively short 500-ms cue-target interval that we used). To address whether the cue-induced modulation of this brain state is related to the cue-induced amplification of target decoding, we correlated each time-frequency sample in Figure 4a with the participant specific magnitude of the main cueing effect on target decoding. Figure 4b shows the resulting correlation map, revealing that those participants who show a stronger alpha attenuation following the cue also have a larger cueing effect on decoding (cluster *p* = 0.024, cluster interval: -20 to 300 ms post target, cluster frequency range: 6 to 11 Hz). This correlation also has a clear posterior topography (inset Fig. 4b).

**Figure 4.**
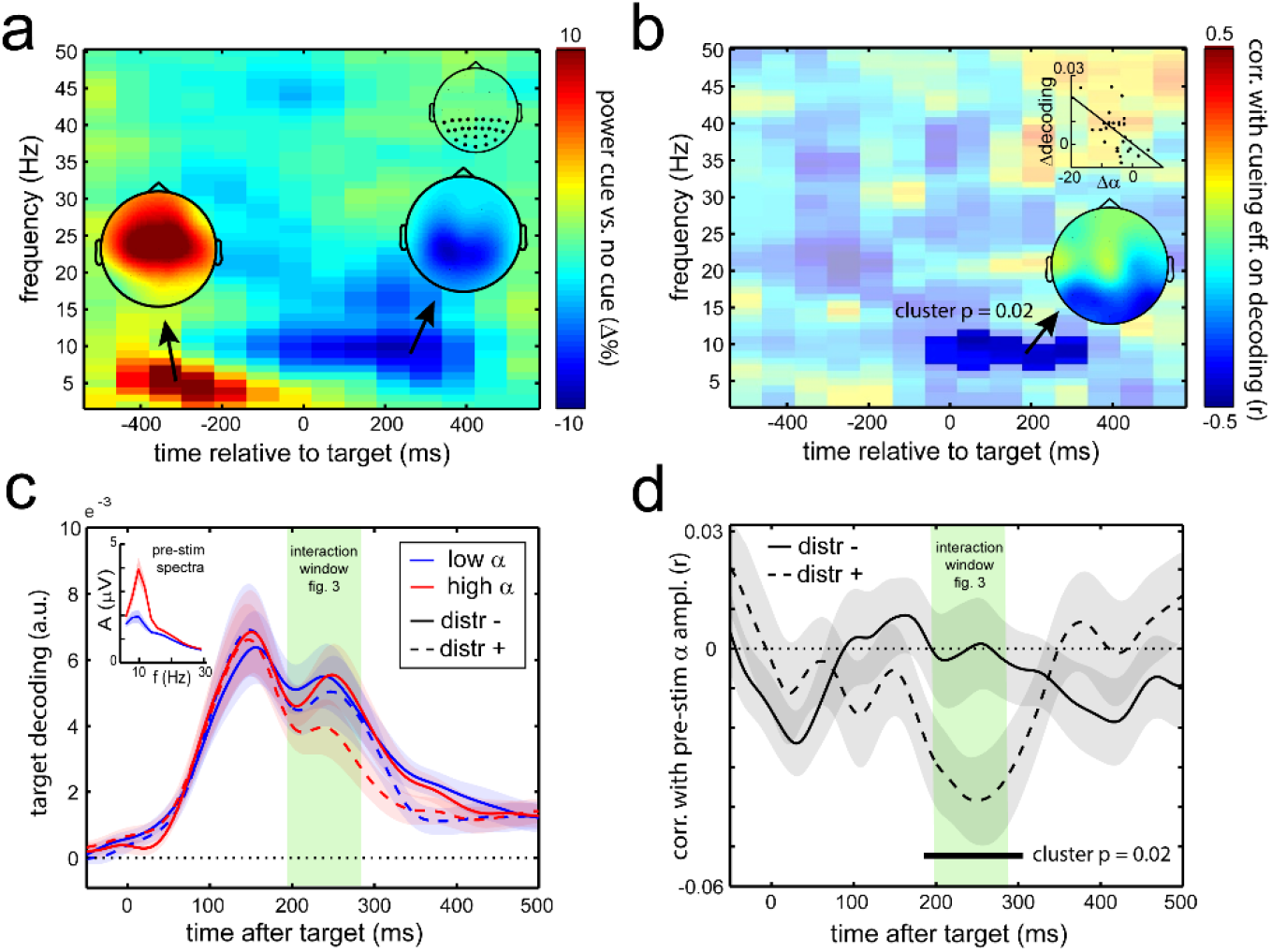
Attenuated posterior alpha oscillations predict enhanced target decoding (across participants) and distractor resistance (across trials). **(a)** Time-frequency plot of the cue-induced modulation in spectral power, expressed as a percentage change (i.e. [[cued – uncued] / [uncued]]*100). Data from all posterior electrodes marked in the inset in the right top. Topographies show modulations from 5-10 Hz in the interval between -400 to -200 ms (left) and from 8-14 Hz in the interval between 0 and 300 ms post-target (right). **(b)** Time-frequency plot of correlation (across participants) of the cue-induced power modulation with the magnitude of the main cueing effect (collapsing distractor present and absent trials) on target decoding (averaged over 118 to 248 ms post-target; see Fig. 3a). The insets show the topography and scatter plot associated with the significant time-frequency cluster. **(c)** Time courses of target decoding in uncued trials as a function of distractor presence and pre-target alpha amplitude (median split). Trials were sorted by alpha amplitude averaged over all posterior channels in the 500 ms pre-target interval. **(d)** Time courses of the trial wise correlation between pre-target alpha amplitude and target decoding, separately for distractor-present and absent trials. Shadings represent ± 1 SE calculated across participants (n = 30). The green shaded band highlights the similarity of the alpha- and distractor-dependent decoding effect with the cue-dependent interaction effect in Figure 3.

We additionally evaluated the relation between alpha states and target decoding across trials. To this end, we focused on all uncued trials (where spontaneous variability is expected to be largest, and where there is no contamination with cue processing) and sorted the trials by alpha amplitude in the 500-ms pre-target window. Figure 4c shows target decoding as a function of both pre-target alpha amplitude (median split; inset for associated spectra) and distractor presence, while Figure 4c quantifies this relation on the basis of a trialwise correlation between alpha amplitude and target decoding. Both analyses yield the same result: the influence of pre-target alpha amplitude appears particularly prominent when distractors were present, whereby attenuated alpha states are associated with a “protective” effect (Fig. 4d, cluster *p* = 0.02, cluster interval: 186 to 307 ms post target). This is highly reminiscent of the interaction effect observed between cue and distractor presence (Fig. 3a). In fact, we noted a strikingly similar time window between both effects (as highlighted in Fig. 4c,d). This analysis thus replicates (using an orthogonal measure) the influence of anticipatory states – either after a cue as in Figure 3, or as reflected in “spontaneously” attenuated alpha oscillations as in Figure 4c,d – on preserving target decoding in the presence of distractors.

We note that these correlations cannot be trivially explained by an increase in signal variance due to higher alpha amplitude. To take away this potential concern, all presented decoding analysis were performed on the time domain signal from which we had removed the 8-14 Hz band using a band-stop filter (and we confirmed that qualitatively similar results were obtained when this filter was not applied).

## Discussion

Combining multivariate decoding and high temporal resolution EEG enabled us to investigate whether and how anticipation influences the amount of sensory information extracted by the brain from target stimuli and temporally adjacent competing distractors. We observed two complementary effects – enhanced target identity coding and delayed interference from temporally adjacent distractors. Enhanced target processing and distractor resistance were furthermore each correlated with alpha oscillatory markers of preparatory attention, thus linking these target decoding effects to two independent operationalisations (cueing and variability in neural dynamics) of “anticipatory state”. The observed effects emerged clearly from a larger set of possible mechanisms by which anticipatory states may help resolve resolve competition between sensory inputs that compete in time.

It is evident that the relevant “coding variable” for perception is not carried by response amplitude per se, but instead by stimulus identity information contained in these responses (see also Kriegeskorte et al., 2006). Our results confirm that anticipatory cues boost the representational quality of visual responses, and reveal that this starts during early sensory processing. Specifically, this “representational boost” peaked around the classical N1 time range. Interestingly, however, while we also observed a parallel cueing effect on ERP amplitude (an amplified N1 response), the magnitude of the cueing effects on target identity decoding and on ERP N1 amplitude were uncorrelated. This suggests that the influence of anticipatory cues on ERP amplitudes and on target identity decoding are mediated by complementary aspects of the EEG (e.g., multivariate patterns that are exclusively available to the decoding analysis), and that the boost in target decoding cannot be simply attributed to a boost in response amplitude (i.e. SNR).

An open question remains what physiological mechanisms may underlie the observed enhancement in target decoding. As likely sources for this modulation, we consider a combination of heightened level of arousal, anticipatory orienting in time (see Nobre et al., 2011), and preparatory upregulation of neuronal populations coding for the relevant feature dimension (i.e. orientation channels; in Garcia et al., 2013 during sustained attention).We speculate that each of these possible “causes” may in turn be mediated by upregulation of the cholinergic system (see Everitt & Robbins, 1997; as well as possibly the norepinephrinic and dopaminergic systems; Warren et al., 2016), in line with the observation that basal forebrain stimulation similarly enhances discriminability of visual input in a rodent model (Goard and Dan, 2009). In the latter work, increased discriminability of visual responses was furthermore linked with decorrelation of neuronal firing rates in visual cortex. It is conceivable that macroscopic states of attenuated alpha oscillations (i.e. alpha “desynchronization”; Pfurtscheller, 1999) provide a non-invasive index of such decorrelated visual activity.

In addition to a direct influence of anticipatory cues on target processing, we also observed a second effect that depended on distractor presence. While distractors always interfered with target decoding, this interference was delayed when targets could be anticipated. Anticipation may therefore enable adaptive perception by allocating a “protective temporal window” from distractor interference (see also, e.g., Shapiro et al., 1997), thereby possibly extending the high fidelity processing of the task-relevant target information and further orthogonalising target and distractor representations. Interestingly, this delayed interference on target decoding by distractors occurred despite the fact that distractor decoding and distractor ERPs themselves appeared not to be delayed. How these observations are to be reconciled remains an important question for future research. One possibility is that, following anticipatory cues, distractor input is being routed to neural populations that show less overlap with those processing targets (despite the fact that both targets and distractors always occupied the same part of visual space). Another possibility is that this reflects increased investment in target processing only until sufficient target orientation information is extracted (after which distractor interference is tolerated again, yielding the observed pattern of delayed distractor interference).

Enhanced target decoding (across participants) and distractor resistance (across trials) were each also related to the attenuation of posterior alpha oscillations – a robust electrophysiological proxy for the level of attentional engagement in human extracranial M/EEG measurements (Foxe et al., 1998; Worden et al., 2000, Thut et al., 2006; Mazaheri and Jensen, 2010; Foxe and Snyder, 2011; van Ede et al., 2012). This was the case both for the task-related modulation by anticipatory cues, as well as for the spontaneous fluctuations in alpha amplitude in the absence of cues. By linking such states to the quality of content-specific early visual brain responses, the current work makes an important extension to a growing body of evidence suggesting a role for such states also in shaping response amplitudes (e.g., Becker et al., 2008), underlying neurophysiology (e.g., Haegens et al., 2011; Snyder et al., 2015), and perceptual as well as mnemonic performance (e.g., van Dijk et al., 2008; van Ede et al., 2012; Myers et al., 2014).

To maximise sensitivity in the current study, we focused on a set-up with simple (but highly effective) temporal warning cues and with large centrally presented high contrast visual gratings. Because of this, we cannot be sure whether our effects are driven primarily by changes in vigilance, voluntary orienting of temporal attention, or both. Still, by linking increased target decoding and distractor resistance observed with cueing also to states of attenuated posterior alpha oscillations, these data do provide a direct link to the voluntary attention literature where such brain states are commonly observed (as discussed above). In future studies, it will be interesting to also track target and distractor identities in relation to more refined attentional and stimulus manipulations (e.g., embedding targets in streams of distractors, cueing different foreperiods, manipulating also spatial and feature-based expectations, etc.). Indeed, as this work showcases, high temporal resolution M/EEG stimulus identity decoding provides a powerful tool for reaching out to previously inaccessible questions regarding cognitive influences on sensory processing in humans (see Garcia et al., 2013; King and Dehaene, 2014; Stokes et al., 2016, for similar arguments). This will prove particularly advantageous in tasks with rapidly changing displays as the decoded output appears largely robust against additive responses (unlike classical ERP responses; compare Fig. 3a with Fig. S1a) while maintaining excellent temporal resolution (unlike fMRI responses).

## Author Contributions

F.v.E, M.G.S and A.C.N designed the study and wrote the paper. F.v.E and S.R.C. conducted the experiments. F.v.E, S.R.C., and M.G.S. analysed the data.

## Acknowledgements

This research was supported by a Newton International Fellowship from The Royal Society and The British Academy (NF140330) as well as Marie Skłodowska-Curie Fellowship from the European Comission (ACCESS2WM) to F.v.E, a Medical Research Council Career Development Award (MR/J009024/1) to M.G.S, a Wellcome Trust Senior Investigator Award (104571/Z/14/Z) to A.C.N, and the National Institute for Health Research Oxford Biomedical Research Centre Programme. The views expressed are those of the authors and not necessarily those of the National Health Service, the National Institute for Health Research or the Department of Health. We also wish to thank Marcel Niklaus and Nick Myers for their input on the experimental design and their assistance during data collection and analysis.

## Methods

Experimental procedures were reviewed and approved by the Central University Research Ethics Committee of the University of Oxford.

### Participants

Thirty healthy human volunteers (10 female; age range 19-35; mean age 25.5 years) participated in the study. All participants had normal or corrected-to-normal vision and either held a university degree or were enrolled in university at time of participation. One participant was left handed. Data from all participants were retained for analysis. All participants provided written informed consent prior to participation and were reimbursed £10/hour.

### Stimuli, procedure, and task

Participants were seated in front of a monitor (100-Hz refresh rate) at a viewing distance of approximately 90 cm. We presented both visual and auditory stimuli (Fig. 1a). Visual grating stimuli consisted of 6 square wave cycles with a total diameter of 18 cm (11.4 degrees visual angle) such that the spatial frequency was approximately 0.53 cycles per degree. We randomly interleaved two types of gratings that were in anti-phase (gratings were either black or white centred), and over which we collapsed in all analyses. Grating orientations were randomly drawn, but were redrawn if within ± 5 degrees from cardinal (0, 90, 180 degrees). We used the same stimuli for target, distractor, and probe displays (see Fig. 1a), although their orientations were independently drawn. Distractors were presented in half the trials and were defined simply by their serial position (i.e. the second grating). Distractors thus acted as visual masks, with the main difference with conventional masks being that distractors consisted of oriented gratings too, enabling us to decode and track both target and distractors identities. Targets and distractors were always presented for 50 ms each, and separated by an inter-stimulus-interval (ISI) of 20, 100 or 200 ms (on distractor present trials). Based on a prior pilot, we anticipated that the 100-ms ISI would yield the largest cueing benefit and we therefore used this ISI in the majority (80%) of distractor-present trials (Fig. 1a). Probe displays always appeared 500 ms after target offset (to avoid response-related contamination of the EEG traces immediately following target onset) and remained on the screen until the participant completed their orientation dial-up using the mouse (or until dial-up time ran out, see below). Auditory cues occurred in half the trials and consisted of 500-Hz pure tones that were presented for 50 ms. Cues indicated that the target would occur after 500 ms, but did not predict whether a distractor would also be present (i.e. cue and distractor presence were manipulated orthogonally). Inter-trial intervals (ITIs), defined as the interval between the response and the next target, did not differ between cue present and absent trials. To maximise the effect of the cues, ITIs were drawn from a truncated negative exponential distribution ranging between 600 and 5000 ms, with a mean of 1000 ms. Because this distribution approximates a flat hazard rate, target onset times were hard to predict, unless a cue was presented.

Participants’ task was to reproduce the perceived orientation of the target stimulus as accurately as possible. To probe perception, we placed a probe grating on the screen, at a randomly drawn orientation. Participants used the computer’s mouse to dial-up the perceived target orientation and clicked once satisfied. Participants were given unlimited time to decide what to report once the probe display appeared, but had to complete their dial-up within 2500 ms once they initiated their response. At response completion, feedback was provided by turning the fixation cross green for 300 ms for responses closer than ± 15 degrees from the actual target orientation (with brighter green colours for more accurate responses); and red otherwise. In total, participants completed 30 blocks of 40 trials each, lasting about 1 hour.

### EEG acquisition and analysis

EEG was acquired with Synamps amplifiers and Neuroscan acquisition software (Compumedics Neuroscan, North Carolina, USA). We used a custom 38-channel set up, sampling all electrodes posterior to the midline from the international 10-10 system and the rest from the associated 10-20 system; thus providing highest density at posterior sites of interest. Data were referenced to the left mastoid during recording, and re-referenced to an average-mastoid reference offline. The ground was placed on the left upper arm. Two bipolar electrode pairs recorded EOG. One pair was placed above and below the left eye (vertical EOG), whereas the other pair was placed lateral of each eye (horizontal EOG). Data were low-pass filtered by an anti-aliasing filter (250 Hz cutoff), digitized at 1000 Hz, and stored for offline analysis. All analyses were run on data with a sampling rate of 1000 Hz. Participant-specific trial-averaged ERP and decoding time courses were subsequently smoothed with a Gaussian kernel with a 15 ms standard deviation.

Data were analysed in Matlab using a combination of FieldTrip (Oostenveld et al., 2011) and custom code. During data preprocessing, we cut out our epochs of interest (relative to target onset), and removed excessively noisy epochs based on visual inspection of the signal’s variance across trials and channels. Artifact rejection was performed on all trials, without knowledge of the conditions to which trials belonged. We additionally removed all trials in which targets and distractors may not have been perceived properly as a result of blinking. To this end, we iteratively removed all trials in which the vertical EOG contained samples with a z-score higher than 5 anywhere within the 400-ms window surrounding target onset.

Time frequency analysis was based on a short-time Fourier transform of Hanning tapered data. We estimated frequencies between 2 and 50 Hz in 1-Hz steps, using a 400-ms sliding time window that was advanced over the data in 80-ms steps.

### EEG orientation decoding

Stimulus orientation decoding was based on the broadband time domain signal that was preprocessed in two ways. First, a 250-ms pre-target baseline was subtracted. Second, the classical alpha band was filtered out of this signal. This was done to ensure that conditional differences in decoding could not be attributed to conditional difference in the signal’s variance related to conditional differences in alpha amplitude (that we anticipated and observed). This is particularly relevant for interpreting the observed correlations of alpha amplitude (across trials) and amplitude modulation (across participants) with target orientation decoding. For filtering, we used an 8-14 Hz band stop Butterworth filter (two pass, filter order 4).

Visual orientation decoding was based on the multivariate (across electrode) Mahalanobis distance metric (as in Wolff et al., 2015; 2017), using the data from the eight most posterior electrodes that showed the largest evoked response (O1, Oz, O2, PO7, PO3, POz, PO4, PO8). For generalization, we applied a leave-one-out procedure. Because this procedure provides a trial-wise decoding estimate, this also enabled subsequent trial wise correlation analyses with pre-target alpha amplitude. Per trial, we calculated the Mahalanobis distance between that trial (the test trial) and all other trials (the training trials) in which the target orientation was at a particular angular difference from the test trial. We did this for 19 bins of training trials centered at angular differences ranging from minus 90 to plus 90 degrees (i.e. in steps of 10 degrees). For each bin, we included training trials whose angular difference from the test trial were within ± 22.5 degrees of the bin’s center. We then mean normalized the resulting distances across all angular bins and averaged the outcome across all trials within each of the experimental conditions. We ran this analysis separately for each time point, thus resulting in a time-resolved orientation tuning profile. For interpretability, we inverted this profile such that angular bins for which neuronal responses that were more similar to the test trial (and thus associated with a *lower* Mahalanobis distance) were associated with *larger* values. To capture orientation decoding in a single metric (per time point), we multiplied the mean normalized tuning profile with an inverted cosine function and summed the result across all angular difference bin (as in Sprague et al., 2016; Wolff et al., 2017). Due to low trial numbers, we did not consider the distractor present trials with an ISI of 20 or 200. Target decoding incorporated all remaining trials, whereas distractor decoding incorporated all remaining distractor present trials.

### Statistical analysis

Behavioural performance data (quantified as the absolute angular deviation between target orientation and reported orientation) were analysed using conventional repeated-measures Analysis of Variance, combined with paired samples t-tests.

Decoding time courses were statistically compared between conditions using cluster-based permutation tests (as described in Maris and Oostenveld, 2007) that effectively deal with the multiple comparisons encountered along the time axes. Specifically, this approach clusters neighbouring samples that survive univariate statistical testing (p<0.05, two-tailed) and evaluates these clusters under a single permutation distribution of the largest cluster that is observed after permuting conditions (at the level of participant specific condition averages). We used 1000 permutations and considered both positive and negative clusters. For target decoding, we evaluated the main effects of cue presence and distractor presence, as well as their interaction (defined as cue present vs. absent for distractor present vs. absent trials). For distractor decoding, we could only quantify the effect of cue presence.

In a complementary analysis, we also evaluated latency differences in these time courses, on the basis of a jackknife approach (as described in Miller et al., 1998). Latency differences were estimated as the temporal difference (between cued and uncued conditions) at which the distractor interference effect first crossed the value associated with 50% of the maximal interference effect (the latter being estimated on the basis of the average of the cued and uncued interference effects). To obtain a jackknife estimate of the reliability of the observed latency difference, we iteratively removed one participant from the participant pool and compared the resulting latency difference to the one observed when all participants were included. The jackknife based estimate of the standard error then allowed us to compare the observed latency difference against 0 (i.e. the null hypothesis of no latency difference) under the student’s t-distribution.

Correlations between EEG amplitudes (across trials) and amplitude modulations (across participants) were quantified using Pearson’s correlation coefficients. EEG amplitudes were averaged over all channels posterior to the midline where alpha amplitude, and its cue-related modulation, were most prominent. For the trial-wise correlation analysis, we partialled out trial number as well as a trial-specific noise estimate that was anticipated to be associated with high amplitude (across most frequencies) and low decoding, thus constituting a potential confounding variable. This noise estimate was obtained by taking the trial-specific variance (across samples) of the high-pass filtered (40 Hz cut-off) data, and averaging this over all posterior electrodes. Correlations were again evaluated using cluster-based permutation analysis to circumvent the multiple comparisons encountered along the time and frequency axes.

All reported inferential statistics involved two-tailed tests, at an alpha level of 0.05.

## Supplemental Information

**Figure S1.**
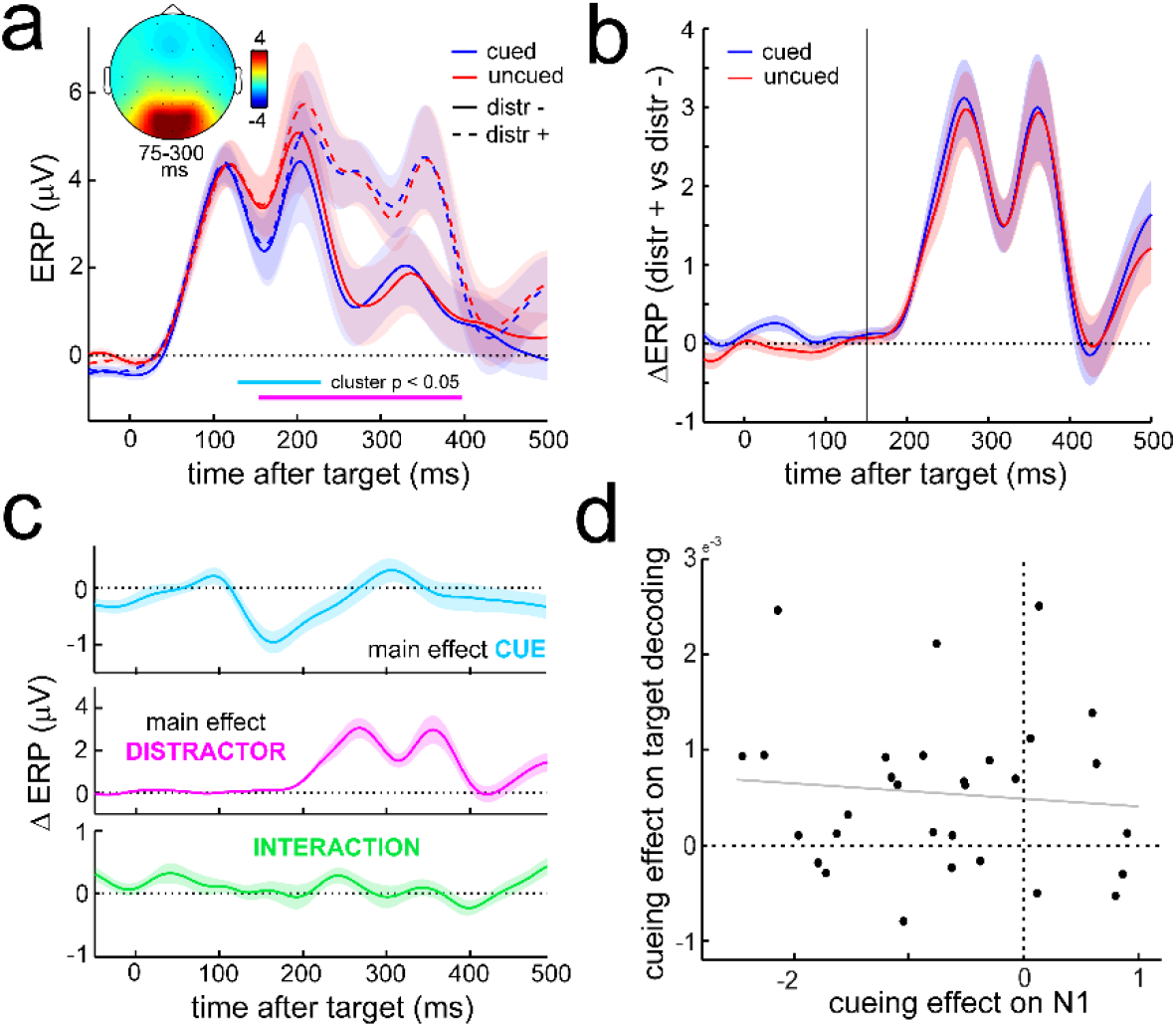
(related to Fig. 3). Event-related potentials as a function of cue and distractor presence. **(a)** Time courses of the ERP in the selected eight most posterior channels as a function of cue and distractor presence (cf. Fig. 3a). Horizontal lines indicate significant temporal clusters for the main effects of cue presence (cyan) and distractor presence (magenta). No significant interaction effect was observed. Data are baseline corrected by a 250-pre-target baseline. Topography inset shows the visually-evoked ERP component between 75-300 ms post-target (collapsed across all trial types). **(b)** Difference ERP for distractor present vs. absent trials, as a function of cue presence. **(c)** Time courses of the main effects of cue and distractor presence, as well as their interaction. The interaction is expressed as the difference between cue present vs. absent trials in distractor present vs. absent trials (cf. Fig 3c). **(d)** Scatter plot showing absence of a correlation between the main cueing effect on the ERP (130-229 ms post-target), and the main cueing effect on target orientation decoding (118 to 248 ms post-target). Individual data points represent individual participants (n = 30).

**Figure S2.**
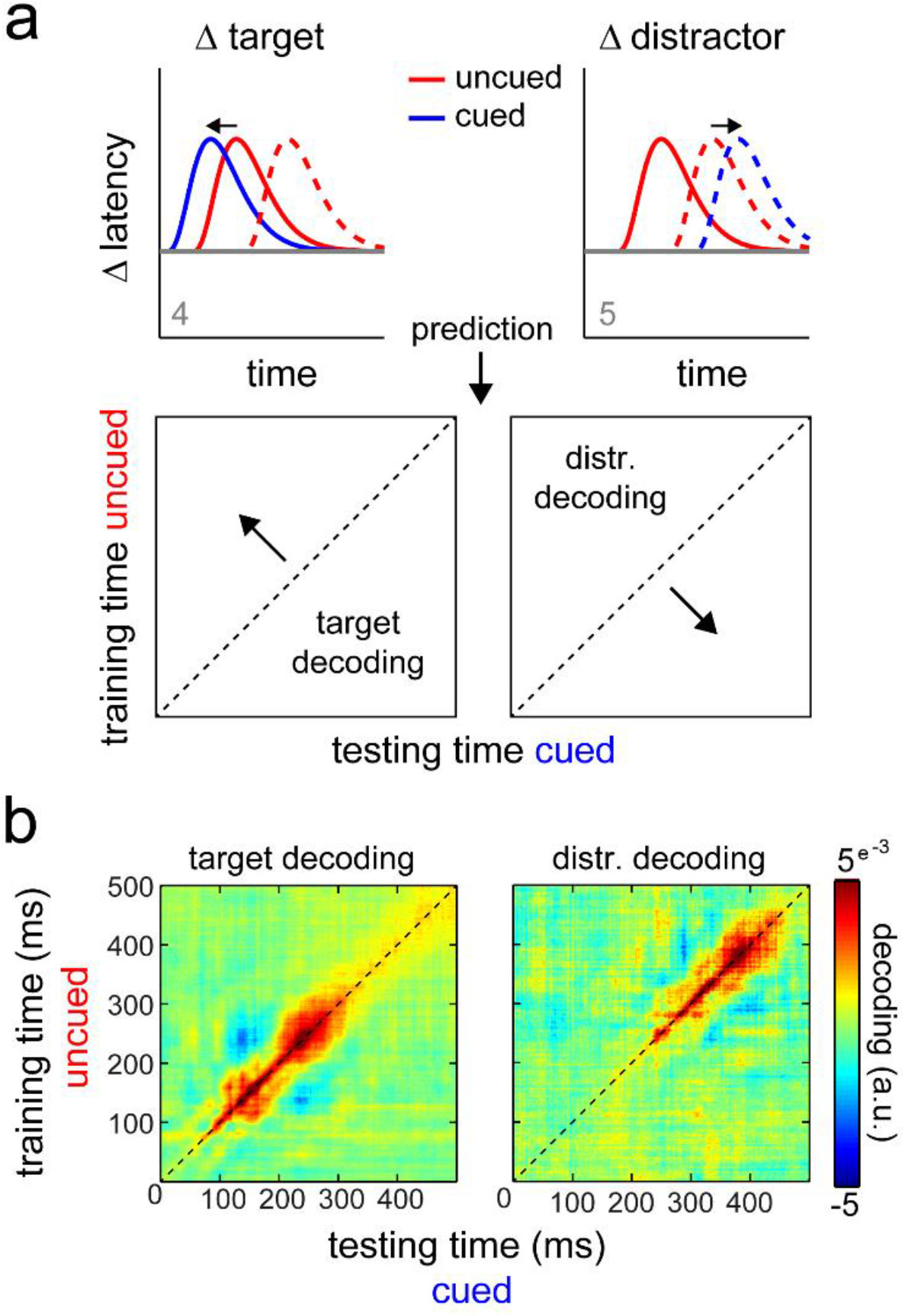
(related for Fig. 2). Cross temporal decoding between cued and uncued trials reveals no cueing effect on the latencies of either target or distractor decoding. **(a)** Schematics of potential cueing effects on target (left) and distractor (right) decoding (taken from Fig. 2), together with predicted pattern in cross-temporal decoding analysis. If cueing results in an earlier target representation, then the cross-temporal plot (training on uncued trials and testing on cued trials) should reveal a leftward shift from diagonal when considering the decoding of the target orientation (e.g., the code at t = 100 during cued trials should resemble the code at t = 100+x in uncued trials). Similarly, if cueing results in delayed distractor coding, then this should reveal a rightward shift from the diagonal when considering the decoding of the distractor orientation. **(b)** Observed cross-temporal decoding data when training on uncued trials and testing on cued trials. For both target and distractor decoding, these plots reveal a clear diagonal focus. Thus, while the “orientation code” is highly dynamic over time, the latencies of this dynamic code remain highly similar between cued and uncued trials.

